# Robotic Assay for Drought (RoAD): An Automated Phenotyping System for Brassinosteroid and Drought Response

**DOI:** 10.1101/2020.06.01.128199

**Authors:** Lirong Xiang, Trevor M. Nolan, Yin Bao, Mitch Elmore, Taylor Tuel, Jingyao Gai, Dylan Shah, Nicole M. Huser, Ashley M. Hurd, Sean A. McLaughlin, Stephen H. Howell, Justin W. Walley, Yanhai Yin, Lie Tang

## Abstract

Brassinosteroids (BRs) are a group of plant steroid hormones involved in regulating growth, development, and stress responses. Many components of the BR pathway have previously been identified and characterized. However, BR phenotyping experiments are typically performed on petri plates and/or in a low-throughput manner. Additionally, the BR pathway has extensive crosstalk with drought responses, but drought experiments are time-consuming and difficult to control. Thus, we developed Robotic Assay for Drought (RoAD) to perform BR and drought response experiments in soil-grown Arabidopsis plants. RoAD is equipped with a bench scale, a precisely controlled watering system, an RGB camera, and a laser profilometer. It performs daily weighing, watering, and imaging tasks and is capable of administering BR response assays by watering plants with Propiconazole (PCZ), a BR biosynthesis inhibitor. We developed image processing algorithms for both plant segmentation and phenotypic trait extraction in order to accurately measure traits in 2-dimensional (2D) and 3-dimensional (3D) spaces including plant surface area, leaf length, and leaf width. We then applied machine learning algorithms that utilized the extracted phenotypic parameters to identify image-derived traits that can distinguish control, drought, and PCZ-treated plants. We carried out PCZ and drought experiments on a set of BR mutants and Arabidopsis accessions with altered BR responses. Finally, we extended the RoAD assays to perform BR response assays using PCZ in *Zea mays* (maize) plants. This study establishes an automated and non-invasive robotic imaging system as a tool to accurately measure morphological and growth-related traits of Arabidopsis and maize plants, providing insights into the BR-mediated control of plant growth and stress responses.

## Introduction

Drought, or limited availability of water, looms as one of the most pressing threats to agriculture. As the world’s population increases, an important challenge is to engineer plants that withstand stresses such as drought while optimizing their growth (Gupta et al., 2020). To realize this goal, we need to understand how plant growth and stress responses are balanced against each other. Such dissection requires precise and comprehensive characterization of growth and drought-related phenotypes and the signaling pathways involved in coordinating growth and stress responses. One such pathway is activated by a group of plant steroid hormones called Brassinosteroids (BRs) that function as critical regulators of plant growth, development, and drought responses (Nolan et al., 2017b).

BRs signal through plasma membrane receptors BRI1 and BAK1 to regulate the activities of BES1 and BZR1 family transcription factors, which control the expression of thousands of genes for various BR responses (Sun et al., 2010; Yu et al., 2011; Nolan et al., 2020). Mutants defective in the BR pathway such as *bri1* are dwarf in stature with reduced stem elongation, shorter and rounder leaves (Li et al., 1996; Szekeres et al., 1996; Clouse et al., 1996), and increased tolerance to stresses such as drought (Northey et al., 2016; Ye et al., 2017; Nolan et al., 2017a; Nolan et al., 2017b). In contrast, gain-of-function mutants in the BR pathway display increased plant growth but often have reduced survival during drought (Ye et al., 2017; Nolan et al., 2017a).

BR phenotyping experiments are typically performed on petri plates at the seedling stage and/or in a low-throughput manner. Since BRs affect plant growth and development at multiple stages of plant life, it would be helpful to comprehensively measure BR related phenotypes in a time-dependent manner with an automated system. Additionally, drought experiments are commonly conducted by subjecting plants to extreme water deficit conditions that are difficult to control or scale to a large number of genotypes. The outcome of drought experiments is often a report on the percent of plants surviving following rewatering, which ignores the dynamics of the drought response and its effect on plant growth (Skirycz et al., 2011). Automated phenotyping of BR and drought responses has great potential to address these short-comings and further define the crosstalk between growth and drought responses.

Recently, several image-based phenotyping systems have been established for large-scale and non-destructive pheno-typing under controlled environments (Bao et al., 2019b; Granier et al., 2004; Skirycz et al., 2011; Serrand et al., 2013; Fujita et al., 2018). Various advanced sensor technologies have been successfully integrated into phenotyping systems, including visible RGB imaging (Minervini et al., 2014; Clauw et al., 2015), chlorophyll fluorescence imaging (Rousseau et al., 2013; Yao et al., 2018), thermal imaging (Zia et al., 2013; Klem et al., 2017), and hyperspectral imaging (Ge et al., 2016; Behmann et al., 2018). While both commercial (Skirycz et al., 2011; Neumann et al., 2015) and custom-built platforms (Apelt et al., 2015; Tisne et al., 2013) have been created, most current systems are limited to two-dimensional (2D) imaging and lack flexibility to administer different types of treatments. In order to more fully understand the relationship between BR-mediated growth and drought responses, we sought to develop a robotic phenotyping platform capable of (1) Conducting time-course observations of plant growth using 2-dimensional (2D) and 3-dimensional (3D) imaging; (2) Administering the BR biosynthesis inhibitor Propiconazole (PCZ) to assess BR response; and (3) Accurately controlling water levels for precise water deficit (drought) treatments.

Here we establish our mobile robotic platform called Robotic Assay for Drought (RoAD) that automates daily weighing, watering, and non-destructive acquisition of 2D and 3D images for BR and drought phenotyping experiments. To make use of the data acquired by RoAD we developed and validated algorithms for automated image processing including rosette and individual leaf segmentation. Subsequent extraction of morphological traits and machine learning approaches allowed us to identify traits that distinguish PCZ or drought treated plants from untreated controls. Using RoAD, we then examined BR and drought phenotypes of Arabidopsis mutants affected by BR signaling, diverse responses of 20 Arabidopsis accessions to PCZ treatment, and BR-mediated changes in the 3D architecture of maize seedlings. Our results demonstrate that RoAD is a valuable tool to study BR-mediated control of plant growth and drought responses.

## Results

### Automated operation of RoAD: a Robotic Assay for Drought

The RoAD system was designed to perform non-destructive imaging, weighing, and watering. The images acquired provide valuable information for measuring the morphological traits of the whole plants as well as the individual leaves. The system consists of a custom-built mobile robot and two tables that can hold up to 240 pots (Figure 1A). The robot was designed as an unmanned ground vehicle (Shah et al., 2016) carrying a Universal Robots UR10 manipulator (Universal Robots, Odense, Denmark) (Figure 1B). An RGB camera (exo249CU3, SVS-Vistek, Germany), a laser profilometer (LJ-V7300, Keyence, Japan), a gripper (2F-85, Robotiq Inc., Canada) and two water drippers mounted on the end-effector of the manipulator. The camera position can be adjusted for plants of various heights. For example, Arabidopsis plants require a lower height than taller maize seedlings. The robot is equipped with a high-precision watering station that is composed of a bench scale (BSQ-0912-001, RMH systems, United States) and two peristaltic pumps (DriveSure, Watson-Marlow, United Kingdom). Two kinds of liquid solutions are available for users to configure different watering regimes. The average error of calculated versus delivered water is 0.38 g (standard deviation: 0.28 g, sample size: 26,078).

**Fig. 1.**
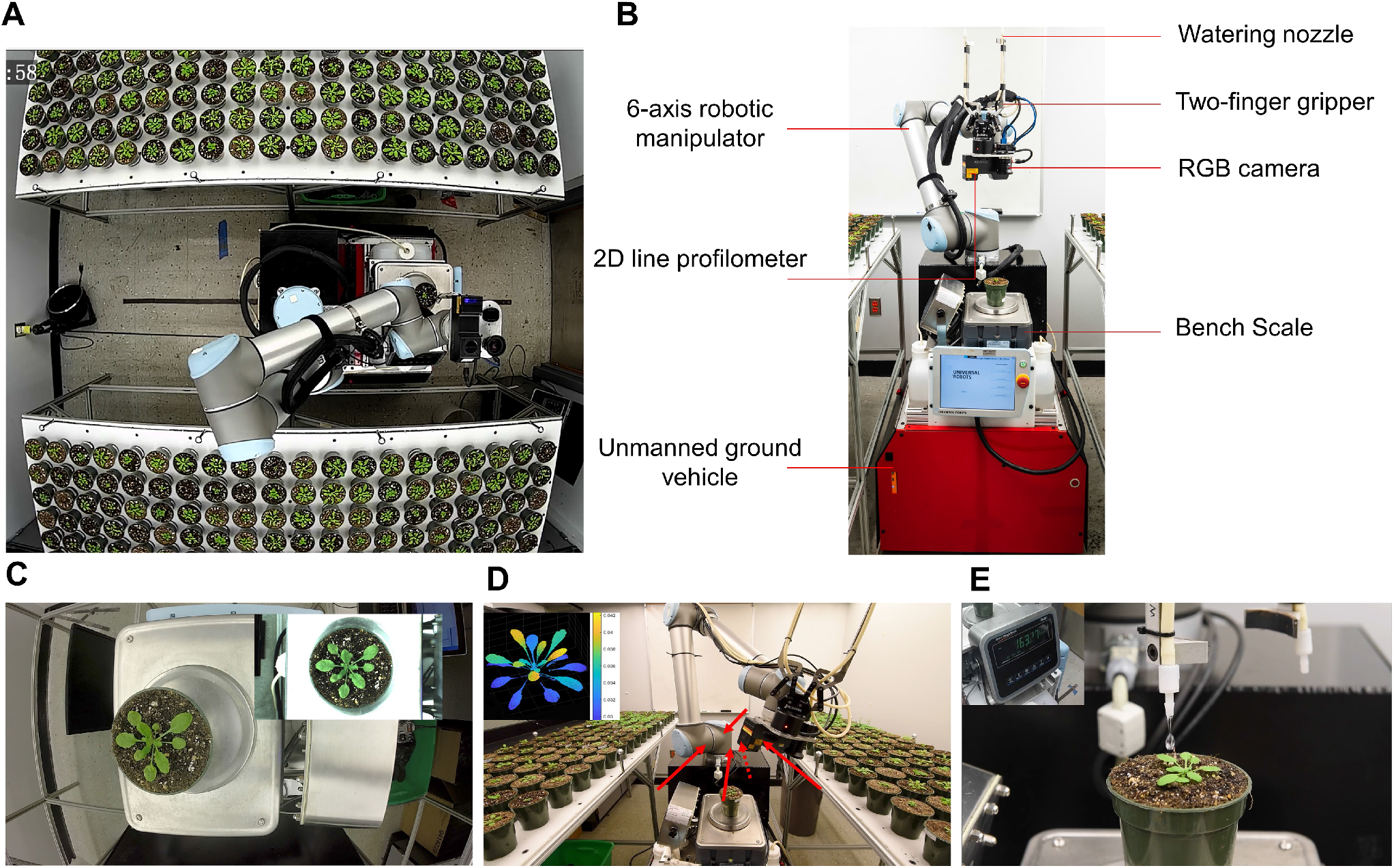
Overview of the RoAD system. (A) Top view of the RoAD system showing the mobile robot and two tables that can hold up to 240 individual pots. (B) The hardware setup of the RoAD system. Instruments for plant handling, imaging and watering are mounted to a six-axis robotic manipulator. RGB camera and 2D line profilometer are used to acquire plant images. The two-finger gripper is used to pick up plants and place them on the bench scale. Watering nozzles attached to the gripper allow for water delivery from two separate water tanks. (C) Top view RGB image of a plant. (D) Multi-view scanning of a plant for the construction of 3D images. Four side-view images and one top-view image are acquired. Arrows indicate the direction of scanning. (E) Example of a plant being watered by the RoAD system. Plant weight is monitored by the bench scale in real-time to allow for precise control of water levels.

An experiment is initialized with a pot map, which stores the attributes of the plants, including plant genotype, replicate, watering solution type and target water level. The plants are imaged, weighed and watered daily. During each data acquisition cycle, the robot parks at one of three positions adjacent to the plant tables. The manipulator is programmed to pick up each pot and place it on the scale. Image acquisition is performed before watering. First, a top-view RGB image is captured (Figure 1C). Then, the plant is scanned from five different perspectives by sweeping the laser profilometer around the plant (Figure 1D) for 3D reconstruction. Multiple scanning perspectives minimize occlusions in 3D reconstruction. The 3D surface model of the plant is reconstructed by cross-registering the 2D RGB image and the 3D point clouds. After image acquisition, the plant is then watered by one of two peristaltic pumps to a predetermined soil moisture level (Figure 1E). Lastly, the pot is transported back to its position on the table. The aforementioned process takes approximately 1.5 minutes. Thus, the full cycle for 240 plants takes around 6 hours, typically yielding 8 gigabytes of raw data (RGB images, 3D point cloud data and pot weight data).

### Automated processing of Arabidopsis plant images

Various strategies for analyzing image data and measuring growth phenotypes have been described (Minervini et al., 2017; Zhou et al., 2017), but general solutions for the segmentation of plants and individual leaves from 3D models remain less developed (Mccormick et al., 2016). The RoAD system provides top-view 2D images of plants and multi-view 3D point clouds. To analyze the large amount of data generated by the RoAD system, we developed a fully automated image processing pipeline.

Our pipeline starts with plant segmentation of the RGB image, followed by segmentation of plants and leaves in the 3D point cloud. Based on the segmentation results, phenotypic trait values are extracted and saved as a csv file for downstream analysis (Figure 2A). The RGB image (Figure 2B) is first converted to a grayscale image (Figure 2C) by Excess Green Index (Hamuda et al., 2016) and then binarized by Otsu’s thresholding method (Figure 2D). Due to the color changes at later growth stages, pot shape information is used to localize regions of interest (ROIs). The Hue channel is then applied to binarize the ROIs and segment drought-treated plants. 3D plant segmentation starts with the points from the segmented RGB image (Figure 2E) projected onto 3D profiles (Figure 2F). The resulting point cloud was refined by Euclidean clustering (Rusu and Cousins, 2011) (Figure 2G). To reconstruct additional details of the plant, the segmented plants from each frame are merged into a single frame (Figure 2H) using an iterative closest point algorithm (Holz et al., 2015). Additional details about image processing are described in the Methods section.

**Fig. 2.**
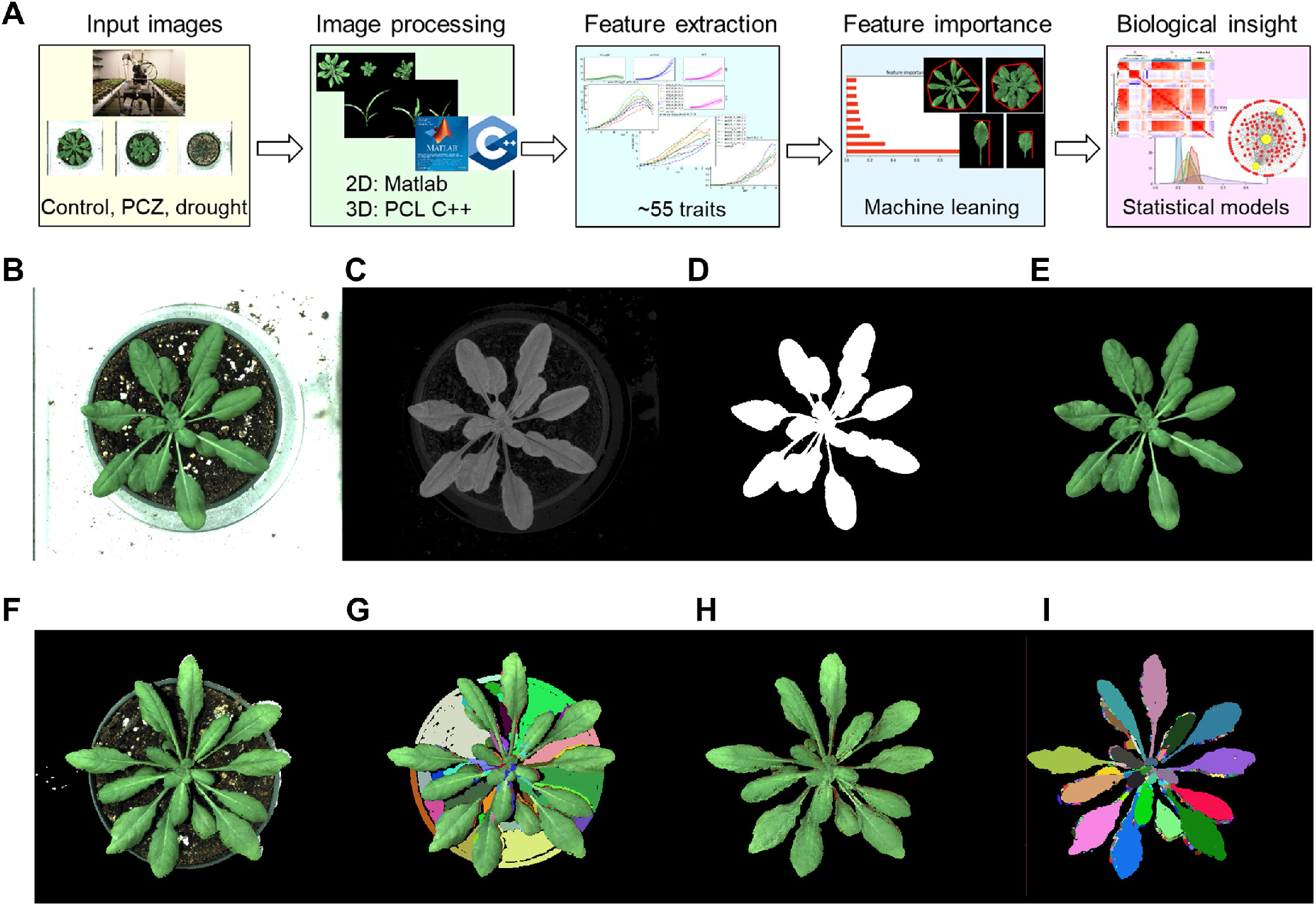
Pipeline for analysis of phenotyping data in the RoAD system. (A) Overview of RoAD system phenotyping and data analysis. 2D and 3D images are captured for plants under control, PCZ, and drought conditions, image analysis is then performed for feature extraction. Fifty-five phenotypic traits calculated from the images are input variables for machine learning classifications. The top features were then used for biological analysis. (B-E) Image processing pipeline for plant segmentation in 2D. The original RGB image (B) is first converted to a gray-scale image (C) by Excess Green Index and then binarized by Otsu’s threshold. After that, the binary mask (D) are used to acquire a plant-only image (E). (F-I) Plant and leaf segmentation in 3D. For each single-view point cloud (F), the segmented 2D plant is projected onto the 3D profile for removing the background, and clustering is used to refine the segmentation (G). The segmented multi-view point clouds are merged to reconstruct the 3D plant (H). Individual leaves are segmented by local features (I).

Previously developed methods for leaf segmentation have implemented deep learning approaches using 2D images (Chen et al., 2019; Liu et al., 2020), and this process requires a large set of manually labeled training data to achieve satisfactory performance. To overcome this challenge, we developed a leaf segmentation algorithm using smoothness and geometrical constraints in the 3D point cloud. A typical Arabidopsis leaf is flat and smooth, and hence the variations between the neighboring surface normals on the same leaf are small. Based on this principle, we applied a normal-based clustering method to the segmented point cloud (Figure 2I). The clustering process begins at the point with minimum curvature value and grows the region by checking local surface normal and point connectivity (Rabbani et al., 2006). Among all the extracted clusters, only the clusters that are sufficiently large and oriented towards the plant center are considered as leaf candidates.

An experiment at full capacity (240 plants) typically lasts 30 days and yields 36,000 images totaling 240 GB of data. Phenotypic information related to plant growth, morphology, and color is obtained from the image data. These phenotypic traits can be categorized into four classes: color-related traits, 2D holistic traits, 3D holistic traits, and individual leaf traits. Color-related traits characterize the color information of the plant, which can be an indicator for assessing plant health (Klukas, 2014). Holistic level traits such as plant area and plant volume assess the entire plant as a single unit. On the other hand, individual leaf traits such as leaf width, leaf length, and leaf angle measure the architecture of individual components. In this study, the three largest leaves from each plant were selected to compute the individual leaf traits. A total of 55 traits were extracted from the segmented images. As expected, we observed a high correlation between 2D, 3D and individual leaf traits that act as proxies of plant growth (Figure 3A). For example, 3D convex area, 2D area and individual leaf area were highly correlated. A full list of the extracted traits and the descriptions of how they were measured can be found in Table S1.

**Fig. 3.**
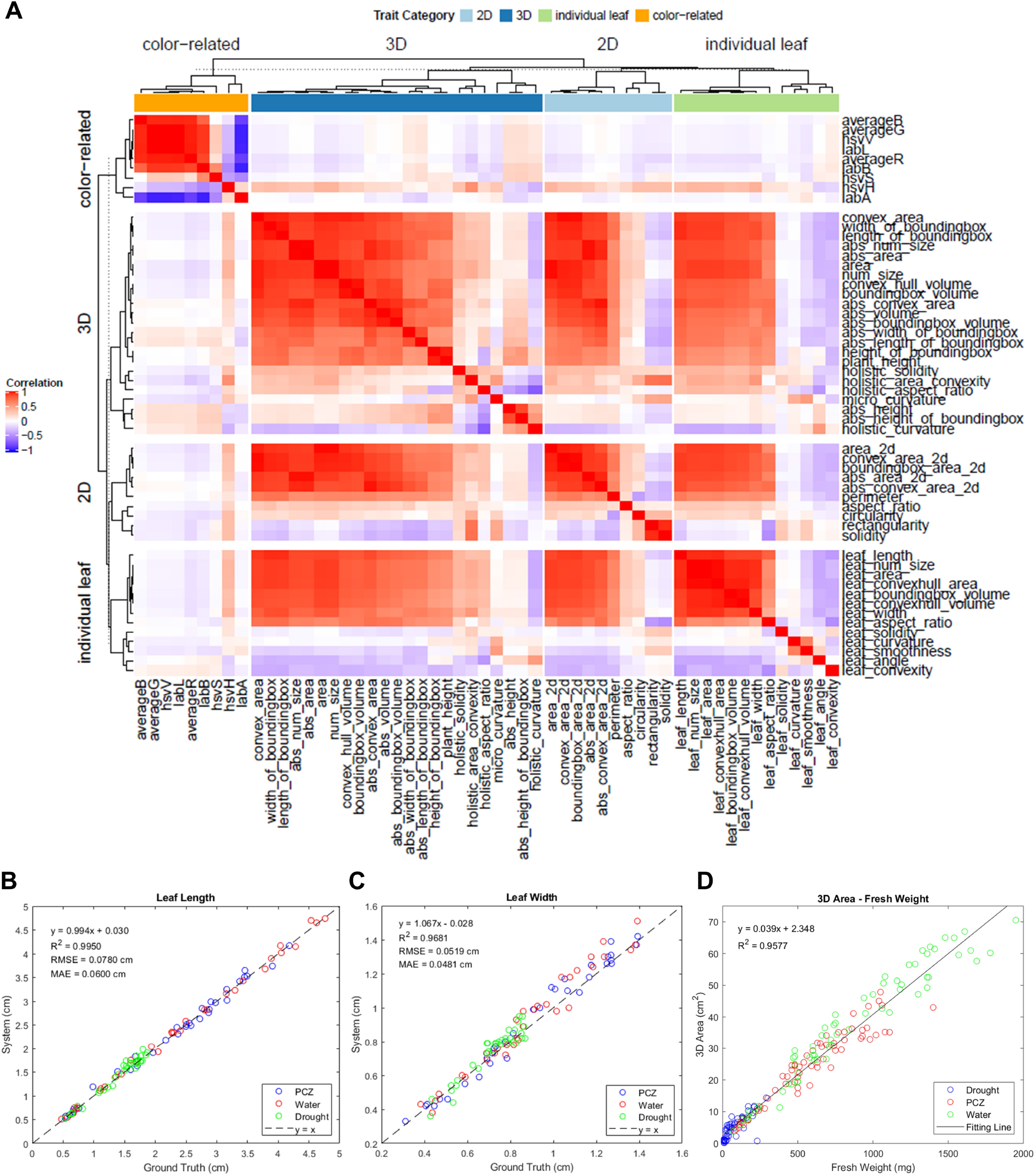
Validation results of the RoAD system. (A) Correlation among the 55 system-derived traits from RoAD image analysis. Traits are grouped by category and ordered by hierarchical clustering. (B) Correlation between system-derived traits and ground truth of leaf length. (C) Correlation between system-derived traits and ground truth of leaf width. (D) Correlation between system-derived 3D area and plant fresh weight.

### Traits quantified by RoAD closely resemble ground truth measurements

To evaluate the performance of the measurements obtained from the RoAD platform and image processing pipeline, 240 Arabidopsis plants of three different treatments were imaged. We also manually collected ground truth measurements of leaf length, leaf width, and fresh weight from the same set of plants. Comparisons between the system-derived traits and manual measurements indicated that the RoAD accurately characterized phenotypic traits of interest (Figure 3B and 3C). For both leaf length and leaf width, the system-derived traits showed high R-squared values and aligned well with the diagonal reference line (x = y), indicating that the RoAD platform has the capacity of measuring leaf traits accurately. We also compared 3D area with plant biomass measured as fresh weight, which showed a strong linear relationship (Figure 3D). Ultimately, the high R-squared values and low mean absolute errors demonstrate the utility of the RoAD platform for automated and reliable measurements of morphological traits.

### RoAD enables BR phenotyping in Arabidopsis

We used the RoAD system to measure growth phenotypes of four Arabidopsis genotypes: wild-type (WT), *bri1-301*, *BRI1P-BRI1OX*, and *bes1-D* under control or 100*μ*M PCZ treated conditions. The seeds of each genotype were germinated in Petri plates for 7 days and a single seedling was transferred to each pot. The plants were allowed to adapt to the soil for 2-3 days before the initiation of a RoAD experiment. During an experiment, each pot started with a well-watered condition. If the gravimetric water content fell below the target level (3g water per g of soil), a specific amount of water or PCZ solution was added to maintain the pot at the desired condition. The 2D and 3D data were collected daily using the RoAD platform. Plants were imaged for 30 days, starting the first day after being set up (DAS). The day when the system was set up was denoted by 0 DAS.

Given the large number of traits reported by the RoAD system, we first asked which of these traits are informative for Brassinosteroid response. We used machine learning to classify WT plants between control and PCZ treated categories. Our analysis attained test accuracies of up to 0.947 (Table S2) and identified a number of traits with high feature importance in distinguishing the BR inhibited (PCZ-treated) plants from the controls (Figure 4A). For example, solidity, which is defined as the ratio of area to convex hull area in 2D, can separate the controls and the PCZ-treated plants effectively (Figure 4B). The solidity of the PCZ-treated plants was higher than that of the controls, indicating PCZ-treated plants show more compact growth. This pattern was also apparent for the holistic area convexity trait (Figure 4C), which is a measure of solidity in 3D, and leaf aspect ratio (Figure 4D), which is a measure of individual leaf shape. PCZ-treated plants showed a higher degree of compactness in 3D models and they have relatively shorter, wider leaves than those of the controls. These macroscopic phenotypic traits observed upon the PCZ treatment are consistent with microscopic ones in which there is a reduction in cell elongation resulting from BR treatment.

**Fig. 4.**
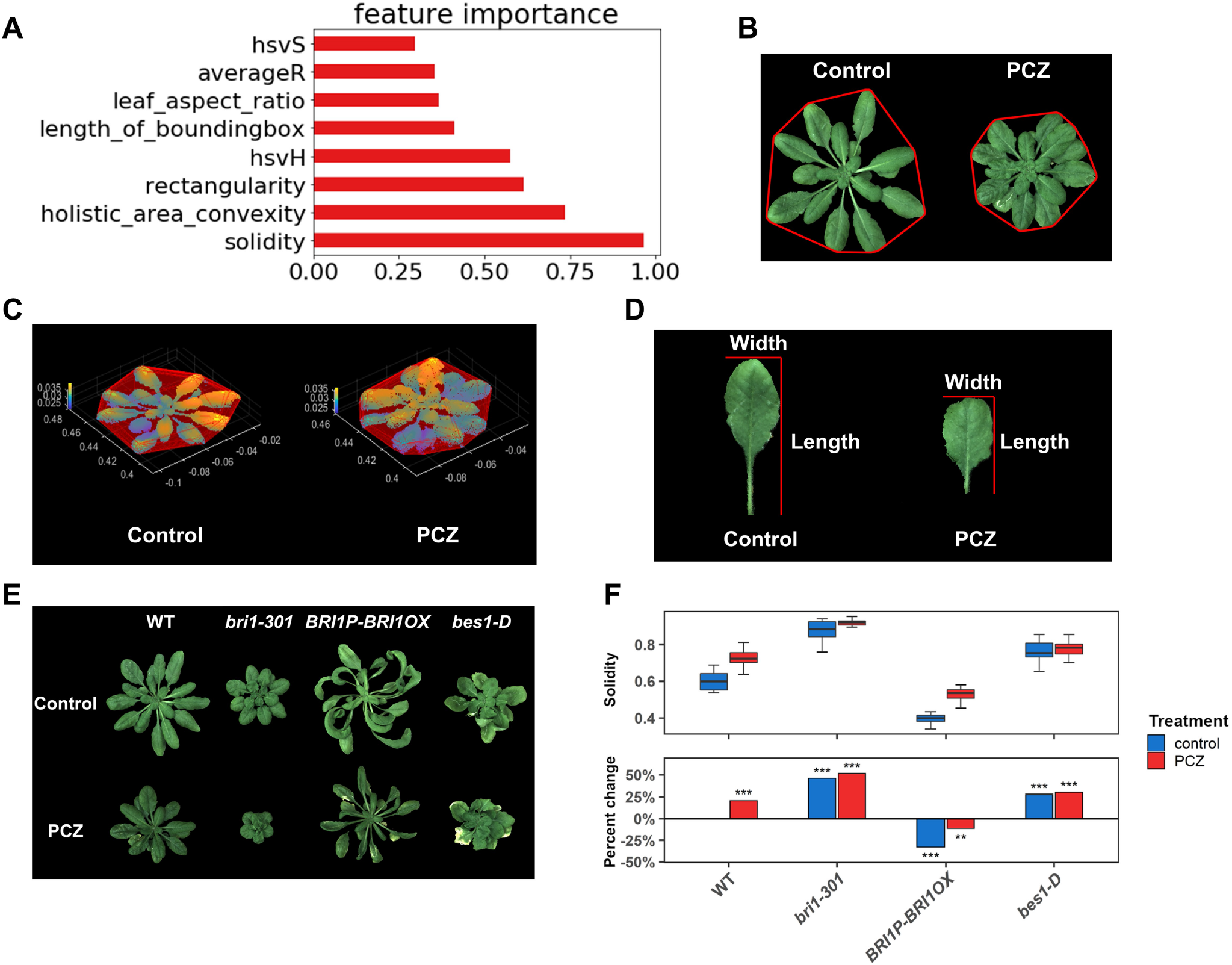
Patterns of BR phenotyping in Arabidopsis. (A) Feature importance from machine learning classification of control versus PCZ-treated plants. (B) Example of a 2D trait: solidity, which is defined as the ratio of projected area to convex hull area. The red outline indicates the convex hull. (C) Example of a 3D trait: holistic area convexity, defined as the ratio of plant area to 3D convex hull area. Plants are pseudocolored based on the depth value and enclosed by 3D convex hull. (D) Example of an individual leaf trait: leaf aspect ratio, defined as the ratio of leaf length to leaf width. (E) Representative images of WT, *bri1-301*, *BRI1P-BRI1OX*, and *bes1-D* plants under control or 100*μ*M PCZ conditions at 30 Days After Setup (DAS). (F) Solidity of WT, *bri1-301*, *BRI1P-BRI1OX*, and *bes1-D* under control and PCZ-treated conditions at 30 DAS. FDR corrected p-values relative to WT control are indicated from a linear mixed model: FDR < 0.001 (***), <0.01 (**), <0.05 (*), <0.1 (+).

If the reduced BR signaling is associated with the more compact growth as measured by increased solidity, then *bri1-301*, a loss-of-function BR receptor mutant (Xu et al., 2008), would be expected to show a pattern similar to PCZ treatment. Indeed, we observed increased solidity of *bri1-301* compared to WT (Figure 4E and 4F). Moreover, solidity showed an opposite trend for *BRI1P-BRI1OX*, which has increased BR signaling (Friedrichsen et al., 2000). However, another gain-of-function BR mutant, *bes1-D* (Yin et al., 2002), did not show increased solidity values (Figure 4F). It is likely that the highly curved leaves of *bes1-D* reduced the rosette compactness due to feedback inhibition of some BR traits in *bes1-D*. Except for *bes1-D*, the order of the solidity of the other three genotypes is *bri1-301* > WT > *BRI1P-BRI1OX*, indicating that increased BR signaling generally reduces plant solidity. A complete list of phenotypic values and corresponding statistical analysis is provided (Tables S3 and S4).

To test how RoAD can be used to phenotype diverse Arabidopsis lines, we examined 20 Arabidopsis accessions from the 1001 genomes collection (Alonso-Blanco et al., 2016; Kawakatsu et al., 2016) under control and PCZ treated conditions (Figure 5A, Tables S5 and S6). These lines were selected due to either increased or decreased hypocotyl elongation in seedlings as assessed by the response to another BR inhibitor, Brassinazole (BRZ) (Asami et al., 2000). We observed concordance between the seedling and plant growth assays in a number of cases. For example, Petergof and Sij 1/96 were stunted in seedling BRZ assays, and similarly they displayed a dwarf phenotype in plant growth assays. Lch-0 had increased growth as seedlings in the presence of the BR inhibitors (Figures S1A-D). Across all 20 lines, there was not a strong correlation between solidity in adult plants in response to PCZ and BL or BRZ responses in seedlings (Figures S1E and S1F). One example is Obh-13, which was resistant to BR inhibition in seedling BRZ assays (Figures S1B-E), but more sensitive to PCZ in terms of solidity in the adult stage (Figures S1A-E). This suggests that additional insight can be gained through BR phenotyping of multiple developmental stages and traits. Consistent with this idea, we found significant genotype by PCZ treatment interactions for 26 traits with 17 accessions having at least one significant difference (FDR <0.1) when compared to the commonly used Col-0 accession (Fig. 5A and B). Interestingly, 3D imaging revealed that Lch-0 plants were taller (Figures 5C and 5D) and had reduced plant area convexity (Figure 5E) compared to Col-0 which coincides with longer hypocotyls in seedling BRZ assays (Figures S1B and S1D). These results indicate that the RoAD system captures traits relevant to BR-regulated plant growth and can reveal additional plant characteristics that might be missed by phenotyping seedlings on petri plates alone.

**Fig. 5.**
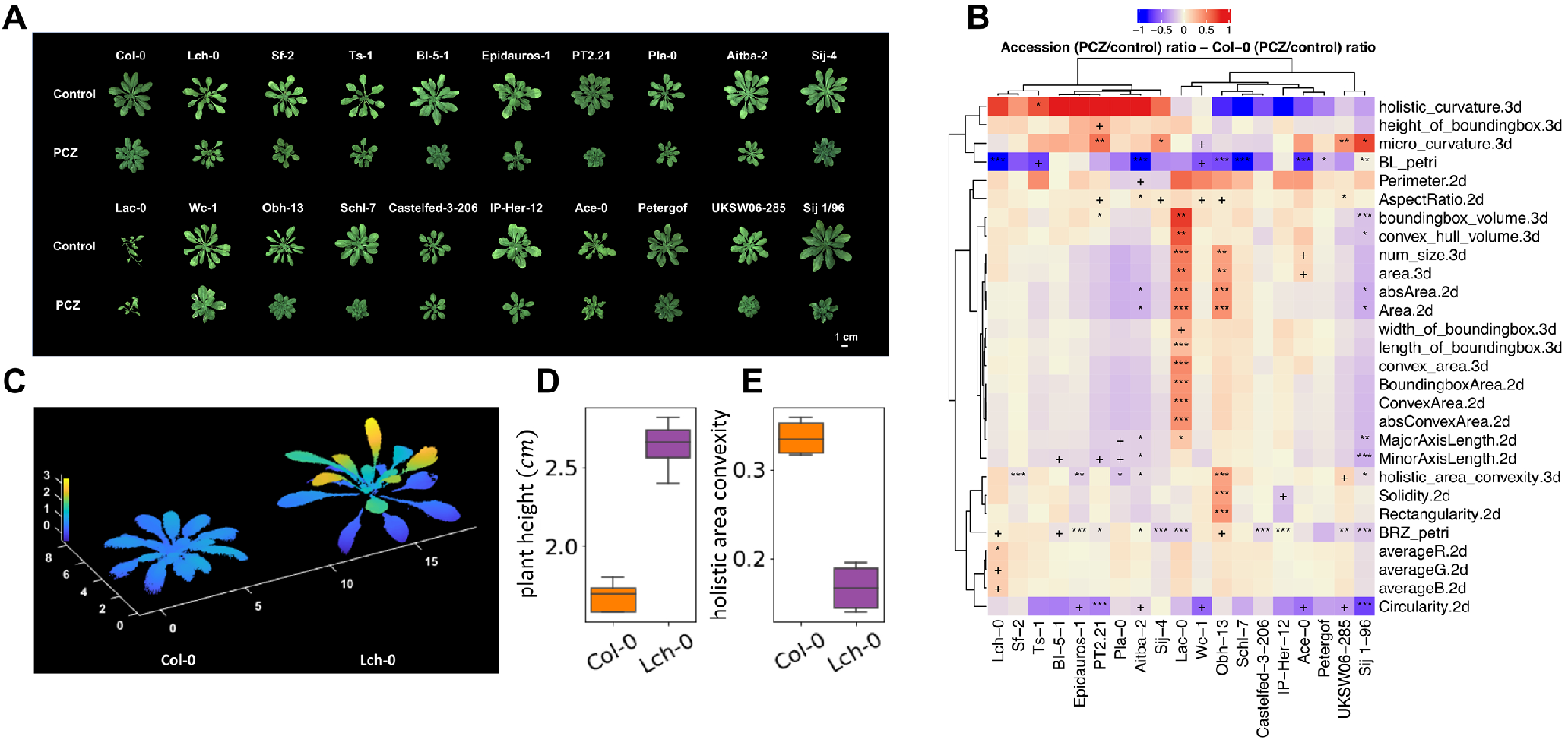
PCZ response varies among Arabidopsis accessions. (A) Representative images of 20 Arabidopsis accessions grown under control or 100*μ*M PCZ conditions on 29 DAS. (B) Heatmap showing genotype by treatment effects for each accession compared to Col-0. BL petri indicates the response to 100nM Brassinolide in light-grown seedlings. BRZ petri indicates the response to 250nM Brassinazole in dark-grown seedlings. All other traits are from PCZ response using RoAD on 29 DAS. FDR corrected p-values are indicated for significant terms from a linear mixed model: FDR < 0.001 (***), <0.01 (**), <0.05 (*), <0.1 (+). (C) 3D models of representative plants of Col-0 and Lch-0 under control conditions on 29 Days After Setup (DAS). (D) Comparison of plant height for Col-0 and Lch-0 under control conditions on 29 DAS. (E) Comparison of holistic area convexity for Col-0 and Lch-0 under control conditions on 29 DAS.

### RoAD precisely controls water levels for drought experiments

The RoAD system can control soil water content for controlled drought experiments in two modes. The first is endpoint drought mode: In this type of experiment, the drought-stressed plants begin the experiment in well-watered conditions and are not watered until they fall below the target moisture level as assessed by gravimetric water content (Figure S2A). One caveat about this method is that drying rates may vary among pots, which has been noted in other automated drought phenotyping systems (Serrand et al., 2013). To address this issue, we implemented a second mode with controlled water deficit ramping. The plants in the drought-stress treatment are kept in well-watered conditions for a set period (e.g. 8 days). Subsequently, the soil moisture level is decreased linearly, enabled by RoAD’s daily weighing and watering regimen (Figures 6A and 6B). In this second mode, both the rate of drying and the timing of water deficit can be more precisely controlled.

**Fig. 6.**
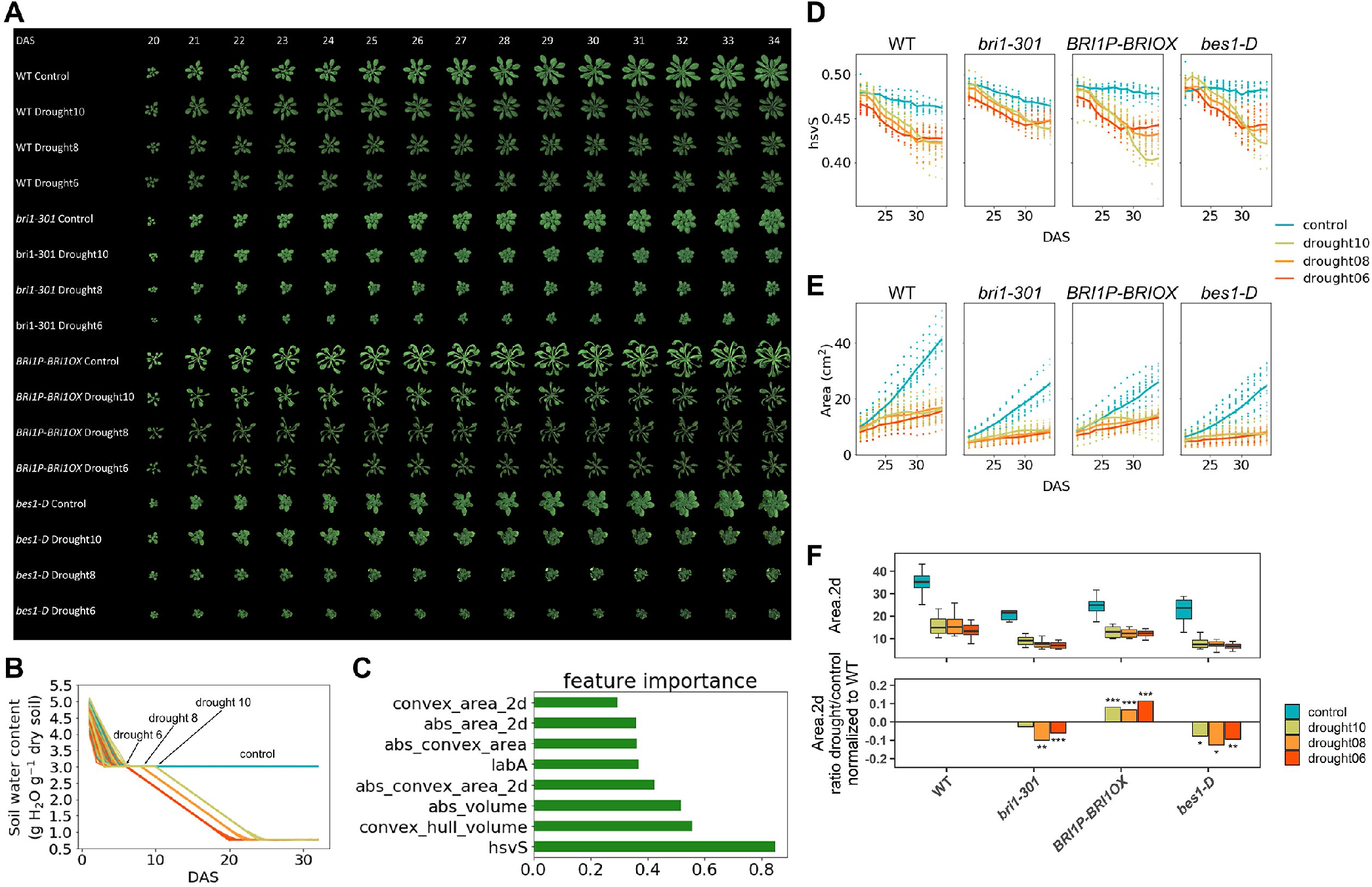
Drought responses in Arabidopsis. (A) Representative images of control and drought-treated WT, *bri1-301*, *BRI1P-BRI1OX*, and *bes1-D* plants from 20 to 34 DAS. (B) Soil water content over time for control and drought-treated plants. For drought treatments the decrease in water levels was initiated at 6 DAS (Drought 6), 8 DAS (Drought 8) or 10 DAS (Drought 10) using controlled water deficit ramping. (C) Feature importance from machine learning classification of WT control and drought-treated plants. (D) Saturation (hsvS) values of WT, *bri1-301*, *BRI1P-BRI1OX*, and *bes1-D* plants under control and drought conditions. (E) Plant area for WT, *bri1-301*, *BRI1P-BRI1OX*, and *bes1-D* under control and drought conditions. Individual plant data are represented by dots, and the group averages are shown with solid lines. (F) Comparison of plant area of WT, *bri1-301*, *BRI1P-BRI1OX*, and *bes1-D* on 32 DAS. FDR corrected p-values are indicated for significant genotype by drought interaction from a linear mixed model: FDR < 0.001 (***), <0.01 (**), <0.05 (*), <0.1 (+).

To establish traits measured by RoAD that are informative for drought phenomics we implemented a similar machine learning classification on WT plants under control versus drought conditions (Table S2). From this analysis, we found that color information could efficiently distinguish control from drought treated plants (Figure 6C). Specifically, drought-stressed plants had lower color saturation values than control plants.

We performed a controlled water deficit ramping drought experiment using WT, *bri1-301*, *BRI1P-BRI1OX*, and *bes1-D* in which water levels were reduced starting at 6 DAS (Drought 6), 8 DAS (Drought 8) or 10 DAS (Drought 10) (Figures 6A and 6B, Tables S7 and S8). We first analyzed the color information but found that the genotypes responded similarly to drought in terms of color saturation (Figure 6D). Next, we examined growth responses in terms of plant area during the drought time series. We observed a more pronounced decrease in growth during drought conditions for both *bri1-301* and *bes1-D* compared to WT (Figures 6E and 6F). These results differ from water-withholding drought survival assays in which *bri1-301* plants have increased survival rates whereas *bes1-D* has decreased survival (Ye et al., 2017; Nolan et al., 2017a). While the conditions for traditional drought survival assays are more severe, the drought conditions applied from RoAD system are milder, which might better represent field conditions. This suggests that monitoring growth during drought using the RoAD system could reveal new aspects of BR-mediated growth and stress coordination. Interestingly, the reduction of growth in *BRI1P-BRI1OX* plants under drought was less severe than that of WT (Figures 6E and 6F). This indicates that some aspects of BR response that are increased in *BRI1P-BRI1OX* may help improve growth under drought. Taken together, the ability of the RoAD system to precisely control soil water conditions and monitor phenotypic traits should prove instrumental in dissecting the cross-talk between BR-mediated growth and drought responses.

### The 3D architecture of BR response in maize seedlings is revealed by RoAD

To extend the RoAD system to a crop plant, we set out to implement RoAD assays in maize. PCZ has been previously demonstrated as an effective BR inhibitor in maize, but the corresponding changes in 3D plant architecture have yet to be explored (Hartwig et al., 2012; Best et al., 2017). To this end, we developed a protocol for RoAD to carry out image acquisition for maize seedling plants nondestructively. During each acquisition cycle, a total of one RGB image and four multi-view point clouds were saved for each maize plant. The multi-view point clouds were filtered and then registered to a single point cloud (Figures 7A). To compute the component phenotypes, a point cloud skeletonization method was used to analyze the maize plant architecture (Figures 7B and 7C) (Bao et al., 2019a; Xiang et al., 2019). A series of morphological traits were automatically extracted. Maize plants were grown in a growth chamber and imaged to evaluate the system and the image analysis algorithm. Comparisons between the measurements indicated that the RoAD platform provides accurate and reliable measurements for seedling maize plants (Figure S3, R2 between 0.93 and 0.99).

**Fig. 7.**
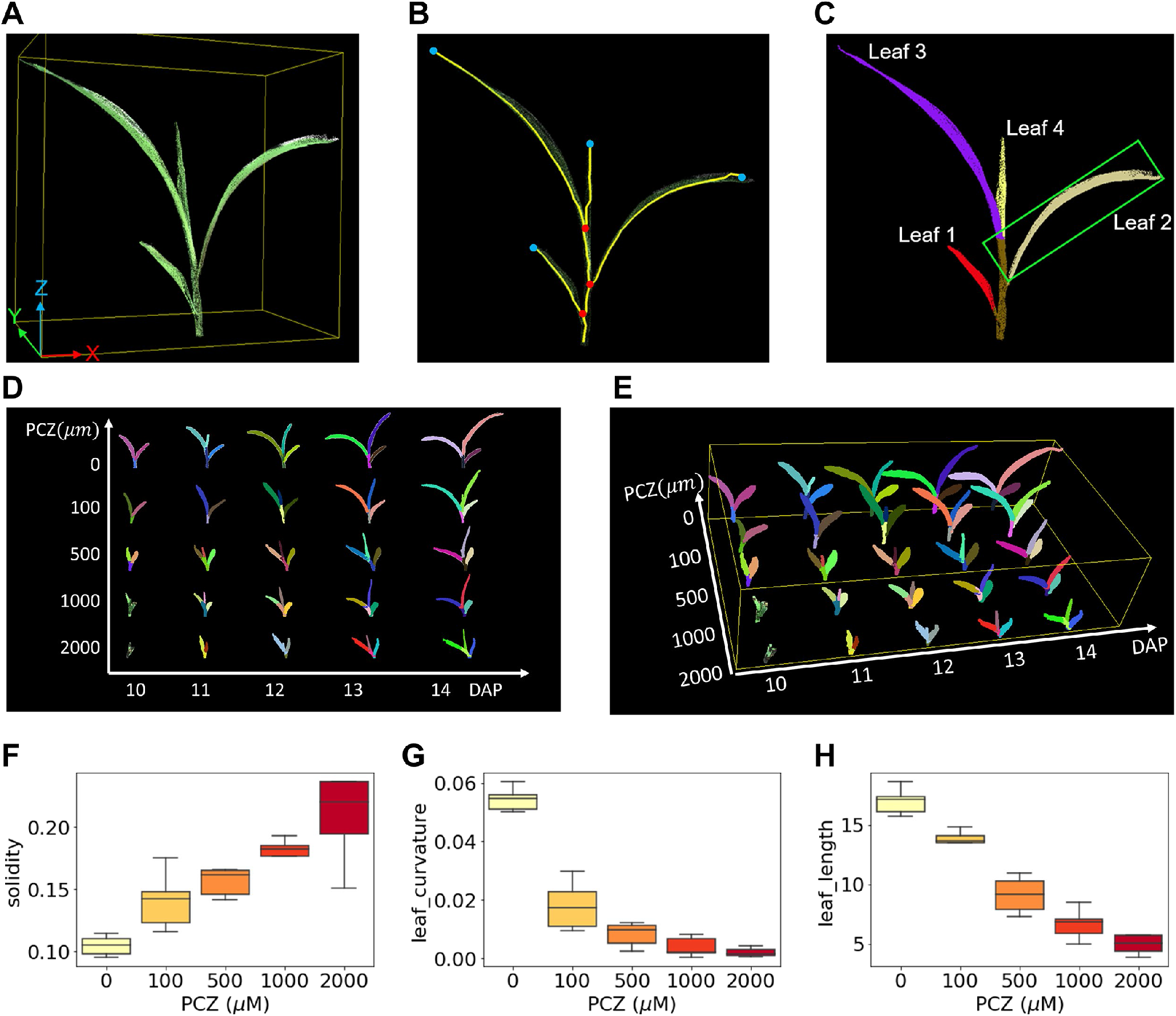
Image processing and BR phenotyping of maize seedlings. (A) 3D point cloud of a maize plant. (B) 3D skeleton of the plant, the blue and red points represent leaf tip and leaf base, respectively. (C) Stem and leaf segmentation. The color indicates individually segmented leaves. (D) 2D view of maize plant growth from 10 to 14 DAP under the indicated control or PCZ-treated conditions. (E) 3D view of Maize plant growth from 10 to 14 DAP under the indicated control or PCZ-treated conditions. (F) Solidity of maize plants under different PCZ levels. (G) Leaf curvature of the second leaf under the indicated control or PCZ-treated conditions.

Thirty maize plants with five l evels o f t reatments (PCZ 0, 100*μ*M, 500*μ*M, 1000*μ*M, 2000*μ*M) were grown to examine the effect of PCZ on maize seedling growth (Figures 7D and 7E, Tables S9 and S10). Images were acquired daily from 10-14 days after planting (DAP). A set of phenotypic traits were extracted automatically using the developed algorithm. We plotted averaged growth curves by plant height, plant width, plant area and plant volume per treatment (Figures S4A-D). PCZ inhibited growth of the maize plants which was evident by the reduction of the plant height, plant width, plant area and plant volume. These effects increased with the PCZ concentration. Consistent with our observations in Arabidopsis, the solidity for PCZ-treated maize plants was also increased compared with controls (Figure 7F). Next, we studied individual leaf traits to gain more detailed insight into the differences observed at the whole plant level. We found that the PCZ-treated plants had lower leaf curvature values than the control plants (Figure 7G). Leaf length also decreased for PCZ-treated plants compared with the control plants (Figure 7H). The decrease in leaf curvature and leaf length at least partially explains the increase in solidity observed and corroborates that PCZ treatment leads to more compact maize seedling phenotypes. The trends observed in our phenotypic characterization of maize seedling PCZ response are congruent with the described roles of BRs in controlling maize growth and development (Hartwig et al., 2011; Hartwig et al., 2012; Kir et al., 2015) and lend new insight into the 3D architecture of this response.

## Discussion

In this paper, we introduce RoAD, an automated phenotyping system designed for BR and drought response in Arabidopsis. The system is capable of watering and maintaining plants at different soil moisture conditions, as well as providing top-view RGB images and 3D multi-view point clouds of plants over time. RoAD incorporates an automatic image processing pipeline, supporting plant and leaf segmentation, and calculation of morphological and color features. The pipeline was validated with manual measurements of plants. Overall, we found that system-derived traits were highly correlated with the ground truth data collected manually. We assessed how traits measured by RoAD vary among BR mutants subjected to PCZ or drought conditions. Additionally, we phenotyped 20 Arabidopsis accessions under control and PCZ treated conditions, which revealed substantial variation in traits affected upon BR inhibition. The system was also used for maize seedling plant phenotyping to demonstrate that it is readily extensible to the analysis of other plant species.

The RoAD system differs from other previously developed phenotyping systems by (1) utilizing a mobile base, which can easily move to and fit in different growth chambers; (2) adopting a six-axis robotic manipulator, making the robot more versatile, dexterous and flexible to acquire multi-view images; and (3) allowing multiple treatments such as PCZ and water limitation. In drought experiments, users can set when water limitation starts, the target water level, and when the target water level is reached. The robotic platform is extendable to other analytical sensors (such as near-infrared, thermal, and probing sensors) and could be integrated into facilities for large-scale plant phenotyping. A limitation of the RoAD system is that during each acquisition cycle, there is a gap of several hours between when the first and last plants of an experiment are processed, which means the timing of imaging and watering varies from plant to plant. To address this issue, we have incorporated a randomized block design that avoids confounding between factors of interest such as genotypes or treatments and the acquisition order. It would be helpful to design multiple robots working in parallel to reduce the time between data collection for different individuals.

Using RoAD and machine learning, we identified solidity as an important feature in distinguishing control from PCZ-treated WT plants. While inhibition of the BR pathway by PCZ and a BR loss-of-function mutant, *bri1-301*, reduced plant solidity, increased BR signaling in *BRI1P-BRI1-OX* decreased solidity (Figure 4). On the other hand, *bes1-D*, which is also a BR gain-of-function mutant, had more complex phenotypes at the whole plant level with increased solidity compared to WT Col-0. It is worth noting that the *bes1-D* mutant used in this study was introgressed into the Col-0 background (Vilarrasa-Blasi et al., 2014), whereas this mutant was originally described in the Enkheim-2 (En2) background (Yin et al., 2002). The increased solidity of the *bes1-D* allele used in this study might be due to highly curled leaves likely due to feedback inhibition on the BR pathway.

By phenotyping 20 Arabidopsis accessions we identified a large array of traits that responded to PCZ treatment differently than Col-0, which is often used as a WT control and reference accession (Figure 5). Additionally, the 3D imaging capabilities of RoAD detected altered plant height in the Lch-0 accession, which would have been difficult to observe from 2D imaging (Figure 5). We noticed that seedling BR response assays did not always correlate with adult plant PCZ response phenotypes, suggestions complementarity among these assays. Our results demonstrate the utility of phenotyping BR-mediated growth responses across different developmental stages, phenotypic traits, and genotypes. Brassinosteroids have extensive cross-talk with drought and several mechanisms impinge on BES1 to balance BR-regulated growth responses with drought survival (Ye et al., 2017; Nolan et al., 2017a; Chen et al., 2017; Xie et al., 2019). Gain-of-function *bes1-D* mutants have reduced survival during drought, whereas loss-of-function *bri1-301* mutants display increased drought survival (Ye et al., 2017; Nolan et al., 2017a). Despite these opposite phenotypes in terms of drought survival, RoAD drought experiments showed that both *bes1-D* and *bri1-301* had more dramatic reductions in growth compared to WT under the drought conditions tested. On the other hand, the growth of BR gain-of-function *BRI1P-BRI1-OX* plants showed less inhibition in response to drought compared to WT. These phenotypes of *BRI1P-BRI1-OX* are interesting in light of the recent findings showing that over-expression of the vascular BR receptor *BRL3*, a homolog of *BRI1*, allows for increased drought survival without compromising plant growth (Fàbregas et al., 2018; Planas-Riverola et al., 2019). Our findings suggest additional complexity in BR-mediated control of drought responses. Future studies should deconvolute the role of various BR signaling components in modulating both growth during drought and plant survival. The precise control of water levels and drought timing enabled by RoAD will enable such investigations.

In conclusion, the RoAD system provides a comprehensive and automated platform for BR and drought response experiments in soil-grown plants. The ability of RoAD to accurately measure morphological and growth-related traits of plants over time and under different treatments should prove a powerful resource to study coordination between BR-mediated growth and stress responses.

## Methods

### Assembly of RoAD System

RoAD consists of a mobile service robot and two elongated tables that support the pots. The robot is made up of an unmanned ground vehicle (UGV), a weighing station, a six-axis manipulator that carries a RGB camera, a laser profilometer, and an electric gripper with two liquid drippers at the fingertips. T he r obot i s a ble t o n avigate i n t he growth chamber and pick up each pot on the tables with high reliability. The base of the UGV has a dimension of 73 cm × 73 cm × 51 cm. The T-slotted aluminum building system (80/20 Inc, United States) was used to build the frame of the vehicle. The UGV is equipped with four mecanum wheels (6” HD, AndyMark, United States) and magnetic guide sensors (MGS1600, Roboteq, United States). The mecanum wheels were driven by four brushless DC motors (BL58-412F-48V GRA60-032, Midwest Motion Products, United States) through two dual-channel motor controllers (FBL2360, Roboteq, United States). The motor controllers, the manipulator and the sensors are controlled by an industrial-grade embedded computer (ML400G-30, Onlogic, United States). To reach all the pots on the tables, the UGV travels along a straight magnetic tape on the floor between the two tables. The UGV can autonomously move between and parked at three positions along the magnetic tape, which are enabled by the magnetic guide sensor and additional magnetic markers next to the magnetic tape. More details about UGV can be found in a published paper (Shah et al., 2016). The two tables (71 cm × 213 cm) are made of rectangular plastic panels and 80/20 aluminum frames. As the magnetic guidance system has a position accuracy of ±1 cm, even spherical metal balls of 2.54 cm in diameter are positioned along the edges of each table for the robot to accurately calibrate its pose with respect to the table. At each workstation, two balls on the near side of the table and one ball on the far side are scanned with the laser profilometer. B all centers are estimated by fitting spheres to the resultant 3D point clouds. Subsequently, the pot positions in a grid system can be located with an accuracy of ±5 millimeters, which is determined by the accuracies of the hand-eye calibration, the synchronization between the laser profilometer and the manipulator, and sphere fitting. The space between adjacent pots on the table allows an approximately 1 cm tolerance. The 3D ball-based pose calibration method is essential to the high reliability of the RoAD platform.

A Graphic User Interface (GUI) was developed to set plant attributes, manage RoAD parameters, and control the RoAD system. To start a new experiment, the user needs to define plant attributes including plant genotype, replicate, watering solution type and target water level. Subsequently, a pot map is generated using a randomized complete block design. In the pot map, each plant has a unique ID. The GUI allows the user to set the drought mode and tune parameters such as the exposure time of the RGB camera, the vertical distance from the camera to the plant, and the speed and acceleration of the robotic manipulator. The user can select various operation modes based on the needs of the experiment. In mode 1, RoAD will grab pots given by the user and put them on the table sequentially. Mode 2 is for daily image acquisition and watering of the plants on the tables. Mode 3 is designed to image and scan a plant that is manually placed on the bench scale.

### Plant materials and growth conditions

Our experiments on the RoAD system included the following Arabidopsis thaliana (Arabidopsis) lines: wild type Col-0 (WT), *bes1-D* (Yin et al., 2002; Vilarrasa-Blasi et al., 2014), *BRI1P-BRI1OX* (Friedrichsen et al., 2000), and *bri1-301* (Xu et al., 2008) and the 20 Arabidopsis accession listed in Table S11. Plants were grown under control (3g water per g dry soil), PCZ (3g water with 100*μ*M PCZ added per g dry soil) or drought conditions (0.75g water per g dry soil). Plant seeds were sown on ½ Linsmair and Skoog plates supplemented with 1% sucrose and stratified at 4°C in darkness for 2-5 days. Plates were then placed in the light at 22°C. After 7 days, the plants were transferred to 10-cm-diameter pots filled with equal weights of soil and soaked in plastic trays with water or PCZ solution. The exact mass of dry soil was determined for each experiment so that gravimetric water content could be calculated to reach the desired soil moisture level. Plants were positioned on the two tables using a randomized complete block design with 4-8 replications per genotype per treatment. Lighting in the growth room was set to a 12-h light and 12-h dark cycle. A dehumidifier was used to maintain the relative humidity at approximately 50%. Weighing and watering were performed once a day for each pot, according to the target conditions. The plants were imaged for 30 days starting from the day when the plants were placed in the phenotyping system. For trait validation, the leaf length and width were measured manually using MATLAB image processing toolbox, which allows determining the distance between pixels. The distance was then converted to an actual length based on the pinhole camera model.

For BRZ response experiments we sterilized seeds for 4 hours in a Nalgene Acrylic Desiccator Cabinet (Fisher Scientific, 08-642-22) by mixing 200mL bleach (8.25% sodium hypochlorite) with 8mL concentrated hydrochloric acid to generate chlorine gas. Seeds were then resuspended using 0.1% agarose solution for plating. Control (BRZ0; DMSO solvent only) or BRZ250 treated (250nM Brassinazole) 1/2 LS plates supplemented with 1% (w/v) sucrose. After seeds were plated, the plates were sealed with breathable tape (3M Micropore) and placed in the dark at 4°C for 5 days. Plates were then exposed to light for 6-8 hours and wrapped in foil for 7 days of growth in the dark. Plates were imaged with an Epson Perfection V600 Flatbed Photo scanner at a resolution of 1200 DPI and hypocotyls were then measured in ImageJ.

BL response experiments were carried out in a similar fashion, except that plates were supplemented with control solvent (BL0, DMSO) or 100nM Brassinolide (BL; Wako chemicals) and plants were grown for 7 days at 22°C under continuous light.

Maize plants were studied to further extend the application of the RoAD system to crop plants. B73 maize seeds were planted in plastic pots in a growth chamber with one seed per pot. The plants were divided into five experimental groups and one control group. The plants in the experimental groups were watered with indicated concentrations of PCZ (100*μ*M, 500*μ*M, 1000*μ*M or 2000*μ*M), and the plants in the control group were grown with water. The plants were cultivated in a growth chamber (16h light/8h dark) with the temperature at 28°C and the relative humidity at 50%. Thirty individual plants were randomly selected from all six groups with five replicates in each group to be inspected by the RoAD system. Image acquisition was performed at five different developmental time points (once a day from 10 to 14 DAP). One RGB image and four multi-view depth images were acquired for each plant. The maize plants were manually transported from the growth chamber to the RoAD system.

### Image processing

Segmentation of drought-stressed Arabidopsis plants in 2D Excess green (ExG) index has been found to be an effective indicator to separate green plants from soil (Hamuda et al., 2016). However, we observed that the plants under water-limited conditions tend to exhibit a dark purple color at late growth stages (Figures 6A and S2A). Accordingly, we implemented hue information to identify the dark purple parts. The pot edges were detected using Circle Hough Transform to aid in the isolation of plants. The part inside the detected circle was considered the region of interest (ROI). The ROI was then transformed to hue saturation value (HSV) color space. An appropriate threshold was then applied to the hue channel of the ROI to separate the drought plant from the soil. The mask images from ExG and HSV color space were combined to acquire a plant-only RGB image. The 2D image processing pipeline was implemented in Matlab R2017a (MathWorks, United States).

Segmentation of Arabidopsis plants in 3D

The processing pipeline of 3D image analysis utilized the Point Cloud Library (Rusu and Cousins, 2011) and the OpenCV library (Bradski and Kaehler, 2008). First, we implemented the Iterative Closest Point algorithm (ICP) to find the global transformation between multi-view point clouds. At this stage, four transformation matrices were obtained, which were later used for merging the multi-view point clouds into a single point cloud. To improve the efficiency of the ICP configuration, the point cloud was down-sampled and filtered to reserve only the parts that were common among the multi-view point clouds (plant, soil and pot). In the next step, we roughly segmented the plant canopy by mapping the fore-ground from the 2D image to the point cloud. However, the resulting point cloud (Figures 2F and 2G) may contain points of partial soil that are occluded by leaves. To resolve this, we adopted Euclidean clustering (Rusu and Cousins, 2011) to filter the point cloud. The point cloud is clustered by Euclidean distance between points (Figure 2G). We found that non-plant clusters, which likely represent soil, were small and close to the pot edge, and the plant clusters were mostly large and have small clusters (young leaves) near the pot center. Therefore, the non-plant clusters are removed by a dynamic size threshold (Eq. (1)) that is a logarithmic function of the point cloud size and the distance from the cluster to the pot center. Cth=SC/Spc*d (1) where SC and Spc repre sent the number of the points in the whole point cloud and the cluster, and d is the distance between the center of the cluster and the center of the pot. Next, the segmented plants from each frame were merged into a single frame using the transformation matrices obtained in the registration process. Finally, in order to remove the duplicate points without losing important information, a voxel grid filter with a 3D box size of 5 mm3 was applied to the combined point cloud.

### Maize plants image analysis and traits validation

A point cloud skeletonization method was introduced to analyze the maize plant architecture and segment individual leaves. The raw data (Figure S5A) were filtered and registered to a single point cloud (Figure S5B). To compute the plant height, the Random Sample Consensus (RANSAC) algorithm (Fischler and Bolles, 1981) was implemented to fit a plane in the merged point cloud to detect the soil (Figure S5C). The points were sliced into layers based on their height and Euclidean clusters were extracted for grouping each layer. The 3D skeleton was generated and mapped to a graph by connecting the centroid of the adjacent Euclidean clusters (Figure S5D). The individual leaf was detected by iteratively traversing the graph from a one-neighbor node (leaf tip, blue points) along a connected path until encountering a three-neighbor node (leaf base, red points) (Figure S5E). Leaves were numbered consecutively, with the first leaf being closest to the soil. The stem was detected as a 3D Hough line (Figure S5F). Based on the segmentation results, a series of morphological traits were automatically extracted (Table S1).

A total of 21 maize plants were grown in a growth chamber and studied to evaluate the system and the proposed algorithm. The plants were sampled at 20 days after planting. The position of the camera was adjusted based on the heights of the plants. After image acquisition, plants were manually measured to collect ground truth data. Plant height and plant width were measured using a ruler. Subsequently, each leaf was cut off to measure the leaf length and leaf area. Leaf length was measured as the distance from leaf base to tip. The leaf was then scanned using an Epson Perfection V600 Flatbed Photo scanner and quantified using Matlab to obtain the area.

### Linear mixed model analysis

A linear mixed-effects model was fit to the trait data using the lme function in the R nlme package (Pinheiro et al., 2020). For each trait and day, the mixed model used raw trait measurements as the dependent variable with fixed effects of genotype, treatment, and their interaction. The random effects structure consisted of a random intercept of plant index within block. Genotype-specific weights were assigned to account for unequal variance across genotypes. The model specification was as follows: lme(raw value genotype * treatment, random = 1|block/index, weights = varIdent(form = 1|geno)). For all plotted data, P values were adjusted for multiple testing according to (Benjamini and Hochberg, 1995).

### Machine learning classification

To understand the underlying relationship between system-derived phenotypic traits and plant responses to PCZ or drought treatments, we constructed two-class classification models based on six machine learning methods: Random Forest, Support Vector Machine, AdaBoost, Gradient Boosting, Extra Trees, and Linear Discriminant Analysis. Feature importance was calculated for each model to assess the relative contribution of each trait in the classification process. The classifications with labeling “1” for control and “2” for PCZ were performed for WT Arabidopsis plants from 15 to 30 DAS. The full dataset of WT plant morphological traits under control and PCZ conditions was shuffled and split into two groups with 70% for training and 30% for testing with 10-fold cross-validation. The performance of the models was evaluated and ranked by test accuracy, and the averaged feature importance ranking was obtained from the top four models. The mean and standard deviation of accuracy, F1, precision, and recall values of the top four classifiers are reported in Table S2. The same processing pipeline was applied to WT plants from 25 to 30 DAS to classify control and drought-stressed Arabidopsis plants. Machine learning classification methods were implemented in Python 2.7.14 (Python Software Foundation, United States) using scikit-image v0.13.0 (Van Der Walt et al., 2014).

## Author Contributions

Conceptualization, T.M.N., L.X., Y.Y., L.T., and S.H.H.; Methodology, L.X., Y.B., M.E., and T.M.N.; Software, L.X., and Y.B.; Investigation, T.M.N., M.E., L.X., N.M.H., A.M.H, and S.A.M.; Resources, Y.B., T.T., J.G., D.S., and L.X.; Data Curation, L.X., T.M.N., and M.E.; Visualization, L.X, T.M.N., and M.E.; Writing - Original Draft, L.X., T.M.N., M.E., and Y.Y.; Writing - Review Editing, L.X., T.M.N., M.E., Y.Y., Y.B., L.T., S.H.H., and J.W.W.; Funding Acquisition, Y.Y., L.T., J.W.W., and S.H.H.

## Acknowledgements

This work was supported by funding from the Plant Science Institute (PSI) at Iowa State University and National Science Foundation (NSF MCB-1181860 to Y.Y. and J.W.W.). We thank Dan Nettleton, Natalie Clark and Hongqing Guo for helpful suggestions, Hao Jiang and Le Wang for technical assistance, and Eugenia Russinova for *BRI1P-BRI1-OX* seeds.

---

**Fig. S1.**
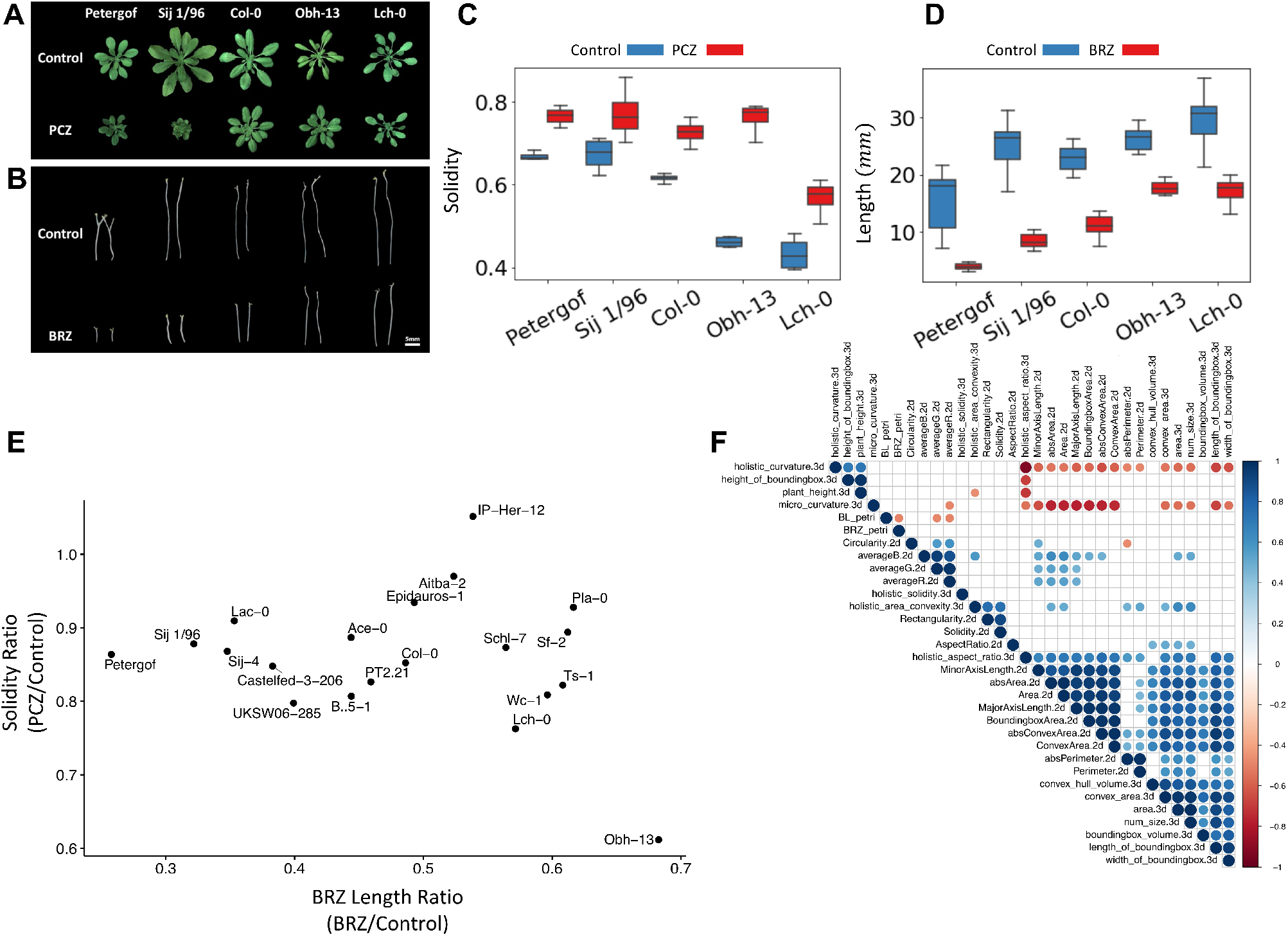
PCZ and BRZ responses of Arabidopsis Accessions. (A) Representative images of 5 selected Arabidopsis accessions grown under control or 100 μM PCZ conditions on 29 DAS. Images are repeated from Figure 5A to for comparison to BRZ phenotypes below. (B) Representative images of 5 selected Arabidopsis accessions grown under control or 250nM BRZ in the dark for 7 days. (C) Solidity of 5 selected Arabidopsis accessions grown under control or 100 μM PCZ conditions on 29 DAS. (D) Hypocotyl lengths of 5 selected Arabidopsis accessions grown under control or 250nM BRZ in the dark for 7 days. (E) Comparison of Solidity and BRZ Length ratios among the 20 accessions phenotyped.

**Fig. S2.**
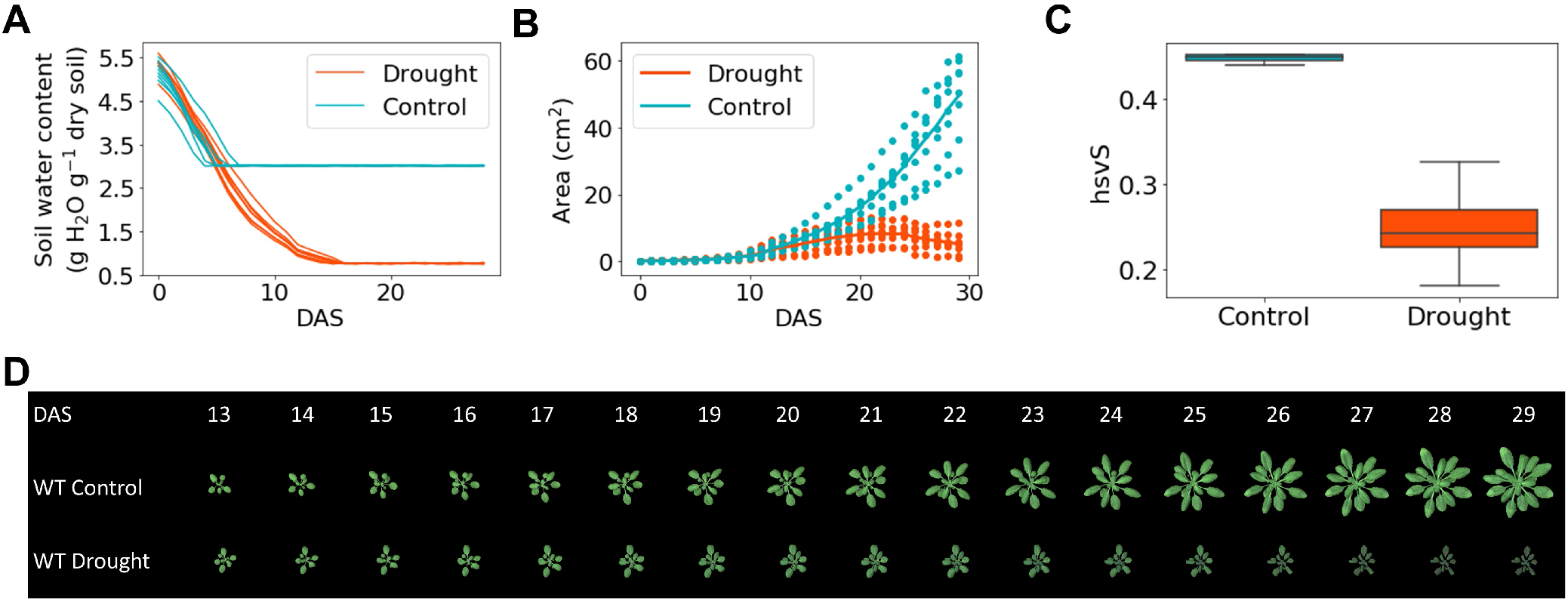
Drought responses in Arabidopsis using RoAD end-point drought mode. (A) Soil water content of control and drought-stressed plants in end-point drought mode. (B) Plant area for WT under control and drought-stressed conditions. (C) Saturation (hsvS) values for WT under control and drought-stressed conditions at 29 DAS. (D) Representative images of control and drought-stressed WT from 13 to 29 DAS in end-point drought mode.

**Fig. S3.**
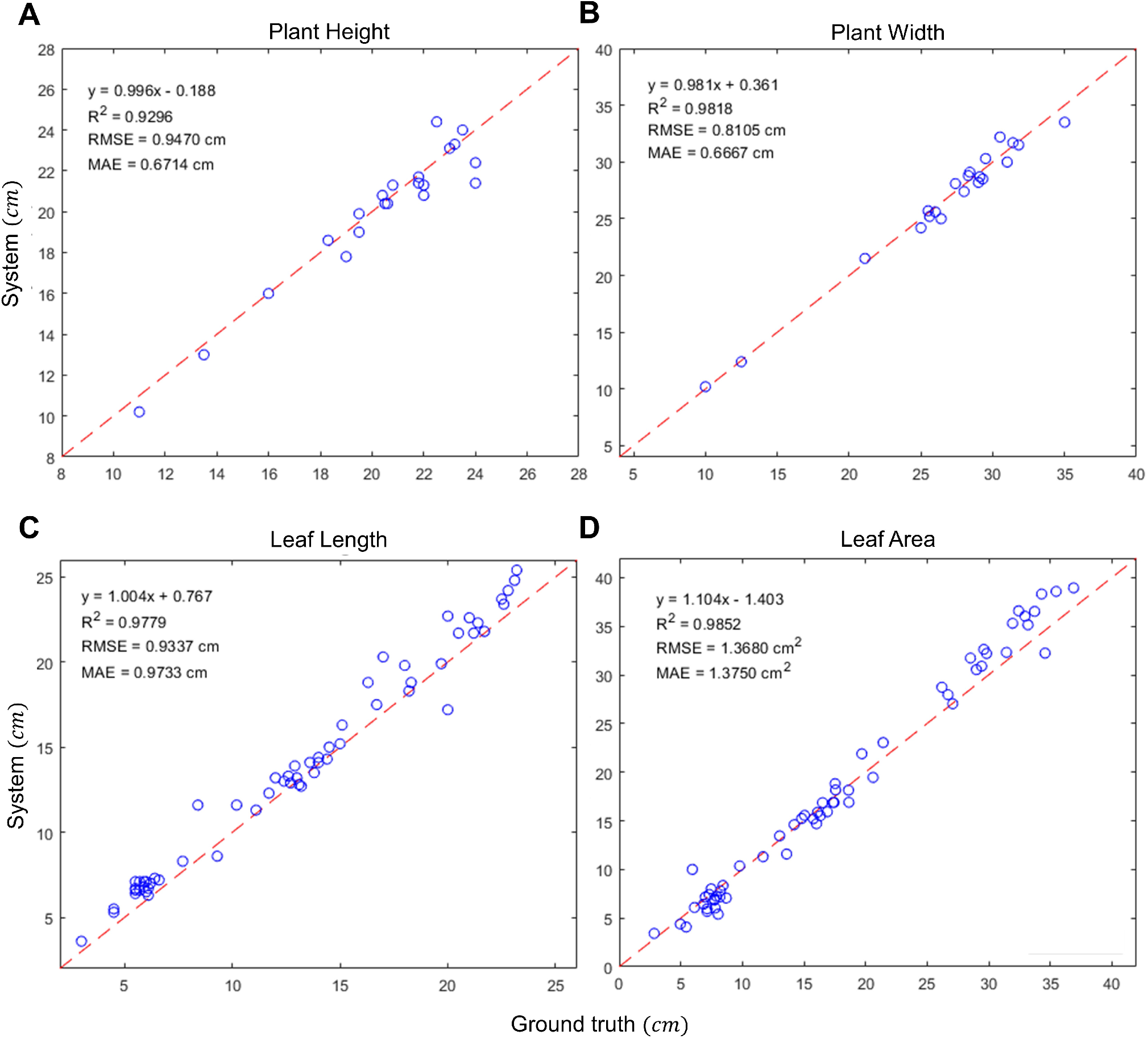
Validation results for maize plants. Linear regression results between the system-derived traits and the manual measurements for plant height (A), plant width (B), leaf length (C) and leaf area (D). RMSE: Root mean square error; MAE: Mean absolute error.

**Fig. S4.**
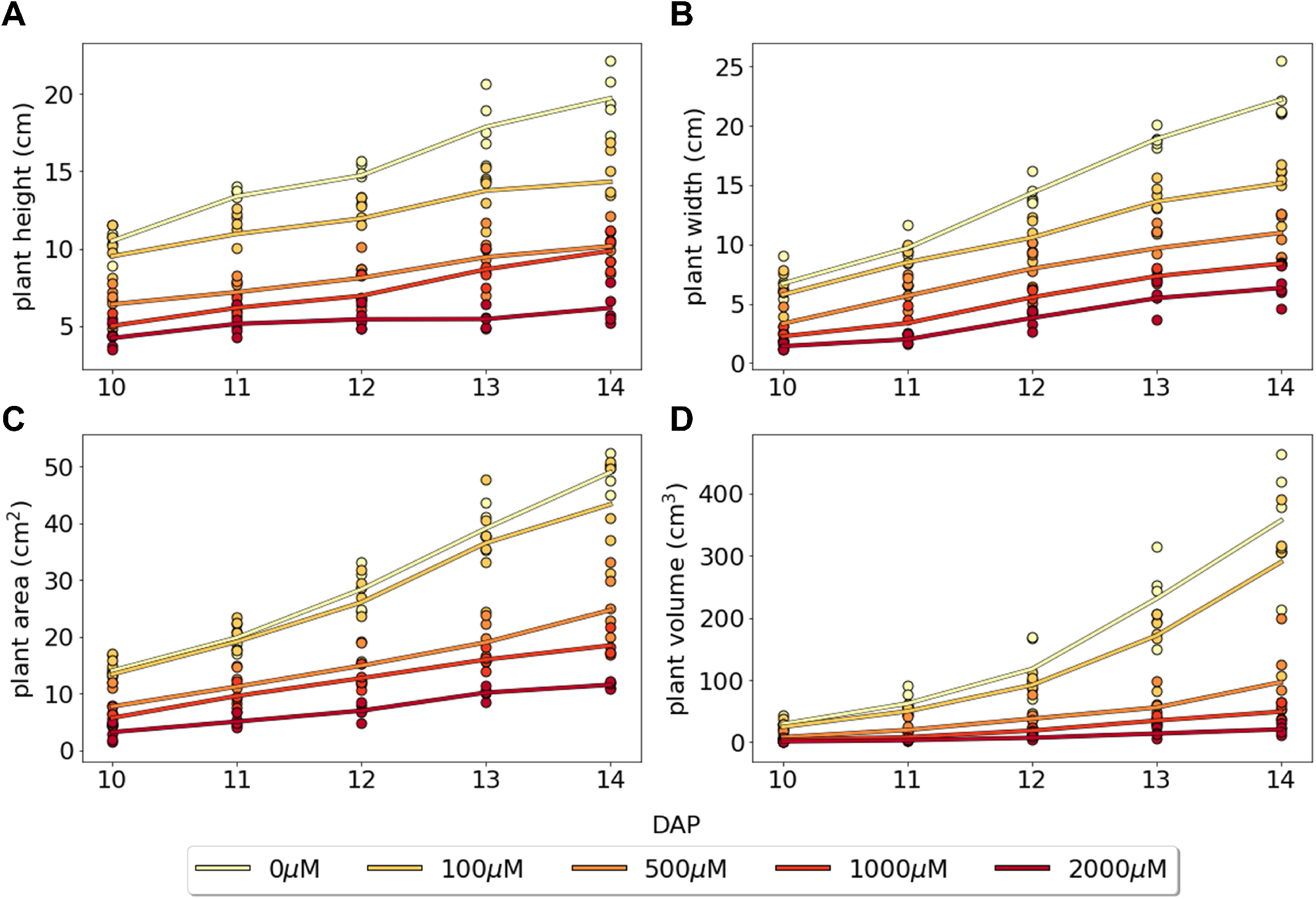
Comparison of traits extracted from 3D point cloud of maize plants. The extracted traits include plant height (A), plant width (B), plant area (C), and plant volume (D). Individual plant data are represented by dots and groups means are depicted in solid lines.

**Fig. S5.**
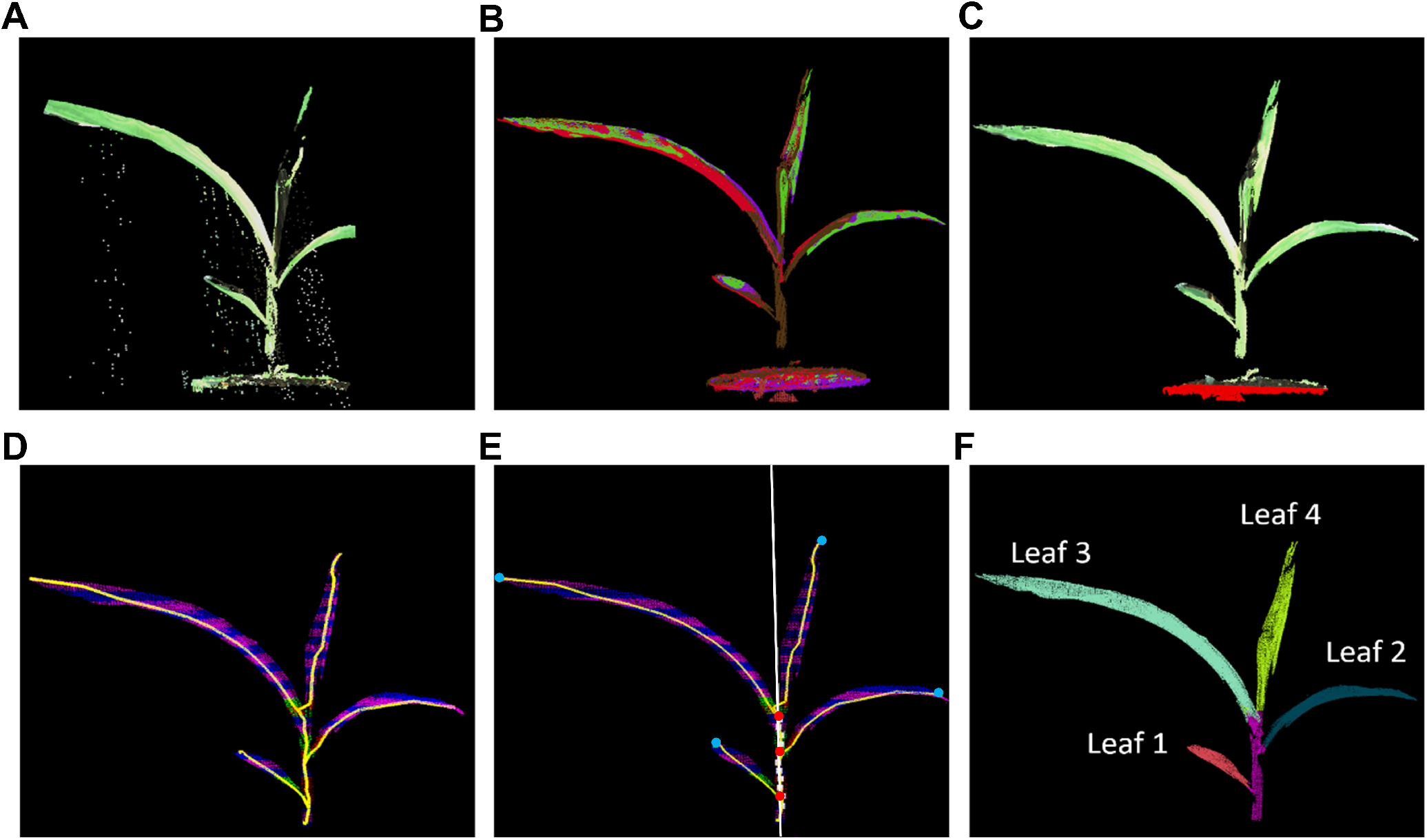
Processing of image data to segment maize seedling stems and leaves. (A) Raw data of a maize plant scanned by the RoAD system. (B) Point clouds sampled from multiple perspectives. (C) Soil detection, red points are soil inliers. (D) Sliced point cloud and the generated 3D skeleton. (E) The detection results of leaf tip (blue points), leaf base (red points), and stem. The white line denotes the stem central line. (F) Segmentation results of the stem and individual leaves.

